# A conserved cold stress syndrome defines insect chill tolerance

**DOI:** 10.64898/2026.10.27.762109

**Authors:** Mads Kuhlmann Andersen, Sebastian F. Jakobsen, Emma Søndergaard Nielsen, Asma Siddika, Clara Garfiel Byrge, Johannes Overgaard

## Abstract

The ability to withstand cold is arguably one of the most important traits defining the biogeographical distribution of insects. For most species, those classified as chill susceptible, cold tolerance is limited by the ability to maintain vital physiological functions at low temperature. However, there is currently no unified understanding of how physiological failure develops during cold stress or how this process explains variation in cold sensitivity within and among species. Although disruption of ion homeostasis has been implicated in chilling injury, it remains unclear whether it represents a common physiological limit across cold stress intensities and degrees of chill tolerance. Here, we tested whether ion balance disruption represents a unifying physiological syndrome of chill susceptibility by simultaneously quantifying cold mortality and hemolymph [K□] in *Drosophila*. We examined this relationship across i) a broad range of cold stress intensities, ii) 25 species spanning extensive variation in cold sensitivity, biogeography and phylogeny, and iii) acclimation groups of *D. melanogaster* developed between 12 and 27°C. Across all comparisons, mortality was associated with the same cold-stress response, characterized by progressive hyperkalemia and mortality following a ∼150% increase in hemolymph [K□] above baseline ([K□]_crit_). In contrast, adaptive and plastic variation in cold sensitivity was explained by differences in the critical temperature at which ion homeostasis failed (T_c_). Extending this framework to published data from other insect orders revealed a remarkably similar [K□]_crit_, suggesting that ionoregulatory collapse represents a conserved physiological constraint defining insect cold survival.

**Summary statement:** Cold tolerance strongly influences where insects can live, yet why species differ so markedly in their ability to withstand cold remains unresolved. Across multiple chill susceptible insects and acclimation groups we demonstrate that insect cold survival is constrained by a remarkably conserved physiological tolerance limit. This limit is defined by an archetypical and uniform disruption of extracellular K^+^ homeostasis where mortality ensues once extracellular [K^+^] has increased by ∼ 150%. Evolutionary and plastic differences in cold resistance therefore reflect variation in the temperature threshold at which ion homeostasis begins to falter rather than divergence in tolerance to the homeostatic disruption. Together, our study provides a unifying overview for variation in cold tolerance across the world’s most diverse animal group.

## Introduction

Most insects have limited capacity for physiological thermoregulation and their body temperature and thermal performance is therefore closely linked to environmental temperature that fluctuates on diurnal and seasonal time scales (1, 2). In accordance, the (in)ability to tolerate thermal extremes is central for the fundamental niche of insect species and metrics of thermal tolerance typically represent powerful predictors of their distribution. This is particularly true regarding cold tolerance limits where physiological resistance often determines species persistence given that acute behavioral avoidance is restricted in the cold, passive and often immobile insects (3–5).

With respect to cold tolerance, insects are typically categorized into tolerance strategies based on whether survival is dictated by the ability to manage the water-ice transition of bodily fluids at very low temperatures or by the physiological consequences of cooling (6, 7). This distinction separates truly cold-hardy species (freeze-avoiding or -tolerant) from “chill-susceptible” species, for which survival is limited by temperature effects on physiological processes rather than ice formation (7, 8). The vast majority of insect species live in tropical and temperate regions and these species typically fall into the category of chill susceptible species which, however, covers a quite broad spectrum of cold sensitivity. Some species succumb at quite mild cold exposure (‘chill sensitive’) while others require long or intense cold exposure before failure ensues (‘chill tolerant’) (9, 10). Furthermore, the species level of chill tolerance is rarely fixed as the level of chill tolerance can exhibit substantial plasticity in response to thermal acclimation (11, 12).

The physiological mechanisms involved in chilling injury has been associated with membrane fluidity changes (13, 14), heat shock protein expression (15), cytoskeletal reorganization (16), immune activity (17), and mitochondrial function (18, 19). However, considerable research has also highlighted that the development of chill injury and mortality is tied to the inability to maintain and regulate extracellular ion homeostasis at low temperature. According to this paradigm, cold injury develops when temperature is reduced below a critical homeostatic threshold (T_c_, **Fig. 1A)** below which active ion- and osmoregulation can no longer mitigate or balance the passive drift of ions and water across cell membranes or epithelia (9, 20, 21). When species are exposed to temperatures below T_c_, the “homeostatic deficit” causes a gradual loss of extracellular ion balance that manifests as hemolymph hyperkalemia (elevated hemolymph [K^+^]) (21). When this hyperkalemic response has reached a critical level ([K^+^]_crit_) it will result in a debilitating level of cellular depolarization that initiates apoptotic and necrotic chill injury (**Fig. 1B**) (22–24). Accordingly, temperatures far below T_c_result in a large homeostatic deficit such that loss of ion balance develops fast and mortality occur within a few hours, whereas stressful cold closer to T_c_ incurs a smaller homeostatic deficit where mortality develops after many hours or days due to a much slower loss of ion balance towards [K^+^]_crit_ (**Fig, 1**).

**Fig. 1.**
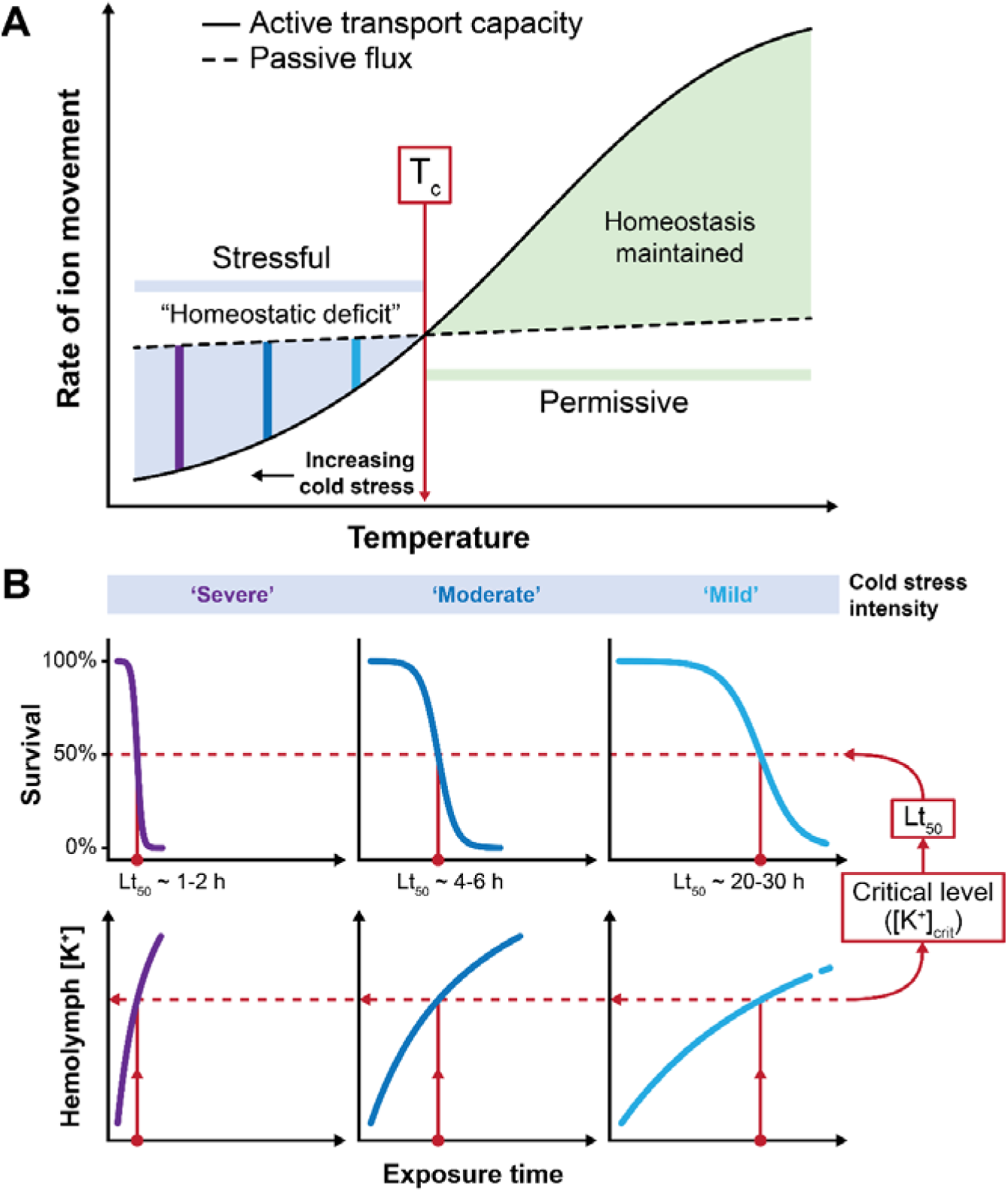
- Conceptual model of the physiological failure underlying cold stress mortality in chill susceptible insects. (**A**) Model of how insect chill tolerance is linked to the balance between active and passive ion transport. According to this paradigm, active transport capacity is much more sensitive to temperature than passive that insects retain surplus capacity for homeostatic ion- and osmoregulation (full line) at permissive temperature (green). However, as temperature crosses a critical temperature (T_c_) and reaches the stressful range (blue) the capacity for homeostatic ion regulation falls beneath the rate needed to balance passive ion flux (dashed line) such that a “homeostatic deficit” manifests, ion balance is gradually disrupted, and cold-related injury begins accumulating. Survival duration at stressful low temperature decreases with decreasing temperature because the intensity of the cold stress (i.e. the homeostatic deficit) increases with lowering of temperature (**B**), however, this model suggests that the proximal cause of cold failure is the same. With the present study, we examine if the archetypical cold stress syndrome defines chill sensitive insects (*Drosophila*) regardless of stress intensity, acclimation status and degree of adaptive cold tolerance among species. Specifically, we hypothesize that mortality (Lt_50_) ensues once a critical level hyperkalemia ([K^+^]_crit_) has developed.

While there is strong evidence that loss of ion balance is associated with cold failure in many insects, the paradigm of a common and uniform cold stress syndrome has not been established across different cold stress intensities. Further, it remains to be determined if chill tolerant species succumb from the same type of physiological failure as their chill sensitive congeners or conspecifics (25, 26) and if differences in tolerance is linked to differences in tolerance towards loss of ion balance (i.e. if [K^+^]_crit_ is higher in tolerant species). In this respect, it has also been suggested that there is a dichotomy of mechanisms involved during acute intense cold stress compared to more moderate chronic cold stress (26, 27). Specifically, acute cold stress has been linked to direct biophysical effects of temperature on macromolecular structure, including reduced membrane fluidity, injurious membrane phase transitions (14, 28, 29) and protein denaturation that could all compromise cellular viability (14, 15, 30). Conversely, prolonged and moderate cold stress is more often associated with the gradual development of homeostatic imbalance and loss of ion balance as explained above (9, 21).

In the present study we leveraged the vast interspecific and plastic variation existing among and within *Drosophila* species to perform a detailed investigation of the link between cold stress mortality and the archetypical ionoregulatory collapse (hyperkalemia) associated with cold stress. We hypothesized that there exists a stress tolerance limit (T_c_) below which injury ensues (**Fig. 1**) and that T_c_ differs systematically among species and acclimation groups. Further, we hypothesized that insect species/acclimation groups are characterized by similar tolerance towards homeostatic perturbation such that mortality is observed once a critical hemolymph [K^+^] is reached ([K^+^]_crit_). We address this question through simultaneous measurements of hemolymph [K^+^] and survival within and among *Drosophila* species to describe the physiological cold stress limit across 1) different intensities of cold stress [from mild to severe, and in split-dose experiments], 2) plastic variation among acclimation groups within a single species (*D. melanogaster*) acclimated to several rearing temperatures, and 3) adaptive variation among species exposed to comparable cold stress using 25 species broadly covering the *Drosophila* phylogeny. Finally, by incorporating published ion balance data from other insect orders in our analysis, we examined whether the physiological stress limit associated with chilling injury is conserved across the broader insect phylogeny.

## Results

To evaluate the relationship between onset of chill mortality and hemolymph hyperkalemia we simultaneously monitored survival (Lt_50_ -time to 50% mortality) and hemolymph [K□] during a range of cold stress experiments. Across all experiments, hemolymph [K□] increased asymptotically towards a new (dis)equilibrium as the cold stress progressed. This allowed us to quantify the magnitude of hyperkalemic response associated with cold mortality (Lt_50_) using an asymptotic model (i.e. [K^+^] at Lt_50_ = [K□]_crit_). This approach was applied across three overarching experiments designed to investigate putative patterns in [K□]_crit_ across *i*) cold stress intensities within three *Drosophila* species, ii) acclimation groups within a single *Drosophila* species, and *iii*) interspecific patterns across 25 *Drosophila* species to investigate phylogenetic and evolutionary patterns. Following the detailed investigation of cold stress tolerance limits ([K□]_crit_), we further assessed cold stress resistance (T_c_) across three species (tropical, temperate, boreal) and across five acclimation groups within one species. This was done by quantifying the lowest temperature (T_c_) that allowed for maintenance of ion homeostasis (i.e. [K^+^] increases at exposure temperature below T_c_ but is stable and regulated above T_c_).

### Effects of cold stress intensity on the progression of hemolymph hyperkalemia

Three *Drosophila* species *(D. montana* [boreal], *D. melanogaster* [temperate], and *D. sulfurigaster* [tropical]) representing a broad spectrum of chill tolerance were each exposed to temperatures constituting mild, moderate, and severe cold stress for each species. In total this resulted in treatment temperatures ranging from -13.5 to +2.5°C across species and treatment intensities (**Fig. 2**). Under benign conditions all three species have hemolymph [K^+^] of ∼ 15 mM and cold stress treatments in all nine experiments (three per species) were associated with highly similar asymptotic increases in [K^+^], reaching [K^+^]_crit_ values ranging from ∼ 38 mM to 46 mM at the time of Lt_50_ (**Fig. 2A**). There were no systematic differences in [K^+^]_crit_ among species (F_2,3_ = 4.4, *P* = 0.127), no difference in [K^+^]_crit_ across stress intensities (i.e. temperatures) (F_1,3_ = 1.7, *P* = 0.282), and no significant interaction (F_2,3_ = 1.3, *P* = 0.289) (**Fig. 2B**). These data suggest, cold stress leads to a similar degree of homeostatic disruption in all three species at Lt_50_, irrespective of whether mortality occurs following acute/intense cold stress (Lt_50_ ∼ 1 h), intermediate cold stress (Lt_50_ ∼ 5 h), or chronic/moderate cold stress (Lt_50_ ∼ 24 h).

**Figure 2.**
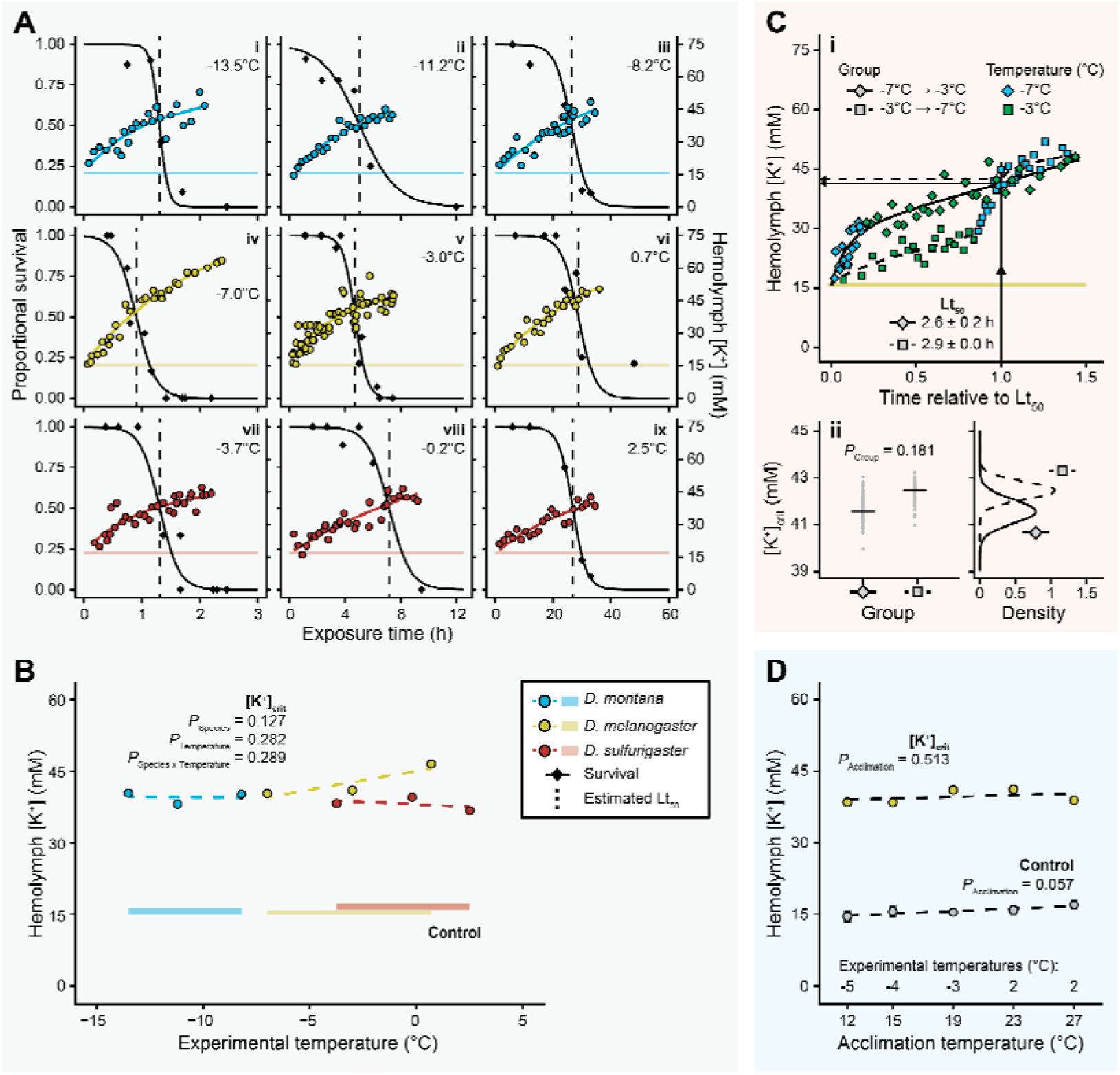
– The critical K^+^ concentration is independent of species-specific cold stress intensities and thermal acclimation but is reached faster at lower temperatures. (**A**) Simultaneous monitoring of survival (black diamonds and line) and hemolymph [K^+^] (circles) in populations of *D. montana* (i-iii, blue), *D. melanogaster* (iv-vi, yellow), and *D. sulfurigaster* (vii-ix, red), representing a gradient of chill tolerant to chill sensitive species. Each species is exposed to three comparable (but species-specific) cold stress intensities and at all conditions we find a similar asymptotic progression of hemolymph hyperkalemia during the cold exposure. Shaded areas indicate control hemolymph [K^+^] and vertical dashed lines the experiment-specific Lt_50_. (**B**) Simulations of [K^+^]_crit_ at time of Lt_50_ across experiments shown in A_i-ix_ reveal that [K^+^]_crit_ is independent of species and cold stress intensity. (**C**) (i) Split-dose experiments where *D. melanogaster* is switched from intense to moderate cold stress and *vice versa* showing that the progression of hyperkalemia follows the stress intensity and (ii) results in similar [K^+^]_crit_ values to those found in static experiments (panel B). (i) Shaded yellow area indicates control values, colors indicate experimental temperature (blue = -7°C, green = -3°C), and specific experiments indicated by the symbols and fitted model lines (diamonds and full = -7°C to -3°C, squares and dashed = -3°C to -7°C). (ii) Small grey points represent [K^+^]_crit_ from each of 100 simulations per group, horizontal lines indicate means, and group-specific data distributions are indicated. (**D**) *D. melanogaster* acclimated to five temperatures and subsequently tested at experimental temperatures invoking similar stress intensities (e.g. Lt_50_ 2-3 hours) revealed that [K^+^]_crit_ was independent of acclimation temperature. Dashed lines indicate trends for control (grey) and critical (yellow) [K^+^] where the slope was not significantly different from 0. Error bars not visible are obscured by the symbols.

To examine if the progression of hemolymph hyperkalemia was additive when stress intensities were combined, we exposed one species (*D. melanogaster*) to sequential exposure to severe (-7°C) and moderate (-3°C) cold stress in a split-dose experiment. Specifically, flies were transferred from one stress intensity to the other upon reaching half of the projected Lt_50_ (based on the experiments in **2A**) (**Fig. 2C_i_**). Regardless of exposure order (intense or moderate stress first) both experiments resulted in similar Lt_50_ values (2.6 or 2.9 h, respectively), and in both groups hyperkalemia progressed according to the cold stress intensity regardless of timing (**Fig. 2C_i_**). As a result, hemolymph [K^+^] reached highly similar [K^+^]_crit_ values (41.6 ± 0.5 mM and 42.5 ± 0.4 mM, respectively; Δ = 0.89 mM, CI = -0.48 to 2.0 mM, Monte Carlo *P* = 0.181) (**Fig. 2C_ii_**). Moreover, the [K^+^]_crit_ from both combinations of split-dose experiments were within the observed range for *D. melanogaster* exposed to constant temperature stress (40.3 – 46.5 mM, **Fig. 2B**) strengthening the observation that [K^+^]_crit_ is independent of cold stress intensity. Accordingly, the cold stress response is similar across the three species and very different stress intensities, the only difference being the rate at which the response develops.

### Acclimation response in D. melanogaster

To test if changes in cold tolerance associated with developmental acclimation changed the relation between Lt_50_ and hyperkalemia ([K^+^]_crit_), we compared the response of *D. melanogaster* acclimated to five temperatures covering almost the full range of temperatures that allows for development in this species (from 12 to 27°C). Developmental acclimation markedly affected cold tolerance and, accordingly, acclimation groups were exposed to different test temperatures in order to impose a comparable stress intensity. Specifically, temperatures ranging from -5 to +2°C were required to achieve similar Lt_50_ values across groups, with Lt_50_ reached after 3.1-4.7 h of exposure (**Fig. S1**). When tested at comparable stress intensity, we observed the same progressive and asymptotic increase in hemolymph [K^+^] for all groups and there were no significant effects of acclimation temperature on [K^+^]_crit_ (mean = 39.6 mM, range = 38.4-41.1 mM; F_1,3_ = 0.5, *P* = 0.513, **Fig. 2D**). Thus, we confirm that developmental acclimation at low temperature increases cold tolerance, but it does not affect the critical level of hyperkalemia tolerated at Lt_50_ (however, see the data on cold stress resistance below).

### Evolutionary and ecological analyses of the hyperkalemic stress syndrome

To broaden the evolutionary investigation of the hyperkalemic cold stress syndrome, we quantified [K^+^]_crit_ in an additional 22 *Drosophila* species (25 total) to examine in detail if there were patterns in [K^+^]_crit_ associated with phylogeny or adaptation to specific environments (**Fig. 3**). The 25 species cover a broad range of chill susceptibility (24 h LT_50_ ranging from -9.0°C to 7.0°C (31)) and include species from the major *Drosophila* subgenera (*Sophophora* and *Drosophila*) associated with several “habitat ecotypes” (31) and latitudinal distributions (see **Table 1** for species information, and **Fig. S2** for raw data). Due to the very broad range of cold tolerance among species, we used different test temperatures for each species, but for all species we aimed to use an exposure temperature resulting in Lt_50_ of ∼ 3 hours (mean ± s.d. = 3.0 ± 1.2 h).

**Figure 3.**
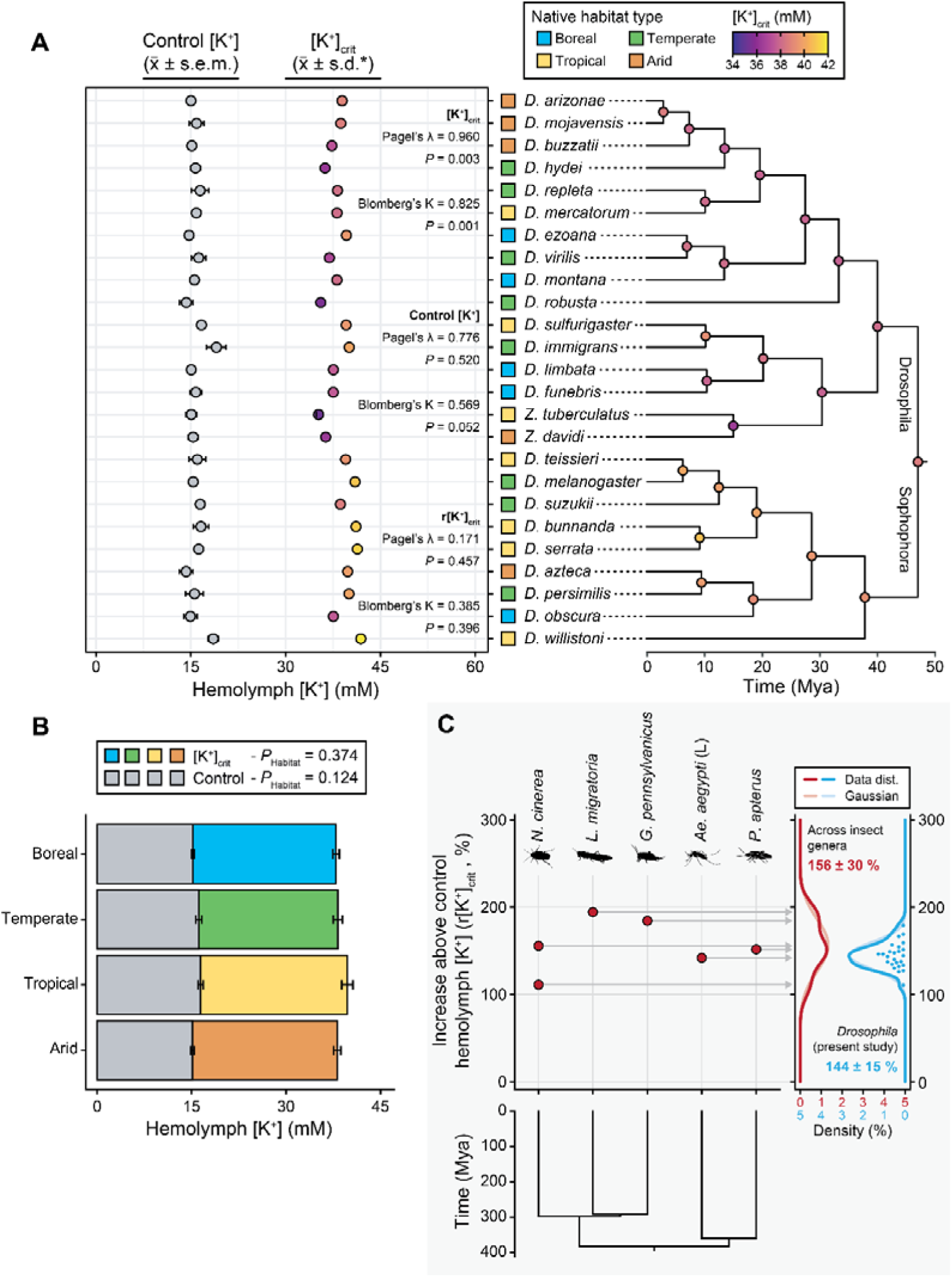
– Multi-species experiment and meta-analysis reveal phylogenetic stability of r[K^+^]_crit_ and suggest an evolutionarily conserved physiological limit to insect cold stress tolerance. (**A**) Control [K^+^] and [K^+^]_crit_ measured across 25 species that broadly cover the *Drosophila* phylogeny. Survival and hemolymph [K^+^] were tracked in experiments designed to constitute moderate cold stress intensity for each species. The resulting [K^+^]_crit_ value for each species (left, colored) is plotted against its position in the time-calibrated phylogeny of the species tested (right), where the evolutionary trajectories of [K^+^]_crit_ are estimated based on maximum likelihood under Brownian motion. Phylogenetic signals (Bloomberg K and Pagel λ) of control [K^+^] and [K^+^]_crit_ are indicated. For each species, the measured control [K^+^] value (grey circles) and native habitat ecotype (squares) are also depicted. Raw data and model fits can be found in **Fig. S2** (**B**) Data from (A) summarized based on habitat ecotype; grey = control [K^+^], color = [K^+^]_crit_. (**C**) Application of the asymptotic [K^+^]_crit_ model to datasets from other species across the insects phylogeny; [K^+^]_crit_ is presented here as a r[K^+^]_crit_ (red points) because insects vary in baseline hemolymph [K^+^]. r[K^+^]_crit_ values obtained from the literature (red) are presented alongside the values obtained for *Drosophila* (blue) and summarized as means ± s.d. Horizontal grey arrows indicate literature data points for [K^+^]_crit_ (relative to baseline) whose data and Gaussian distribution are indicated by opaque and translucent red lines, respectively. *Drosophila* r[K^+^]_crit_ values are indicated by blue points and the data and Gaussian distributions by opaque and translucent blue lines. Error bars not visible in panel A are obscured by the symbols. *This s.d. is based on the Monte Carlo resampling simulation values for [K^+^]_crit_.

**Table 1.** – Information on species used in the present study. Species are described based on their subgenus (*D* = Drosophila, *Z* = Zaprionus, *S* = Sophophora), phylogenetic subgroup, native habitat type, and cold tolerance (critical thermal minimum, CT_min_; low temperature leading to 50% population mortality, 24 h LT_50_). Information on stock origin is also indicated (NDSSC = National *Drosophila* Species Stock Center, Cornell University, NY, USA), and the collection site if available.

| **Species (subgenus)** | **Subgroup** | **Native habitat\*** | **CT<sub>min</sub> (°C)\*** | **24 h LT<sub>50</sub> (°C)\*\*** | **Stock origin** | **Collection** |
|---|---|---|---|---|---|---|
| *Drosophila arizonae (D)* | *repleta* | Arid | 1.9 | -5.0 | Longdon lab | Unknown (received 2020) |
| *Drosophila mojavensis (D)* | *repleta* | Arid | 1.1 | -5.4 | NDSSC | Anza Borrego Desert, California (2014) |
| *Drosophila buzzatii (D)* | *repleta* | Arid | 3.5 | -4.9 | NDSSC | Sardinia (2006) |
| *Drosophila hydei (D)* | *repleta* | Temperate | 2.1 | -3.0 | Kristensen lab | Ehime, Japan (2016) |
| *Drosophila repleta (D)* | *repleta* | Temperate | 4.5 | -3.0 | NDSSC | Izaleo, El Salvador (1953) |
| *Drosophila mercatorum (D)* | *repleta* | Tropical | 6.2 | -1.2 | Kyoto Stock Center | Unknown (received 2020) |
| *Drosophila ezoana (D)* | *virilis* | Boreal | -2.4 | -8.5 | Hoikkala lab | Finland (2019) |
| *Drosophila virilis (D)* | *virilis* | Temperate | 0.8 | -7.1 | Kyoto Stock Center | Japan (<2012, 1987) |
| *Drosophila montana (D)* | *virilis* | Boreal | -3.1 | -9.0 | Hoikkala lab | Finland (2019) |
| *Drosophila robusta (D)* | *robusta* | Temperate | 1.5 | -7.2 | NDSSC | Rocky Point, New York, USA (2004) |
| *Drosophila sulfurigaster (D)* | *immigrans* | Tropical | 6.9 | 7.0 | Kellermann lab | Cardwell, Australia (2016) |
| *Drosophila immigrans (D)* | *immigrans* | Temperate | 6.1 | 1.0 | Sørensen lab | Odder, Denmark (2020) |
| *Drosophila limbata (D)* | *funebris* | Boreal | 4.5 | -8.4 | Lankinen lab | Lahti, Finland (2014) |
| *Drosophila funebris (D)* | *funebris* | Boreal | 1.4 | -6.6 | NDSSC | Tehran, Iran (unknown) |
| *Zaprionus tuberculatus (Z)* | *armatus* | Tropical | 4.1 | -0.9 | Longdon lab | Unknown (received 2020) |
| *Zaprionus davidi (Z)* | *armatus* | Arid | 5.4 | -1.0 | Longdon lab | Unknown (received 2020) |
| *Drosophila teissieri (S)* | *melanogaster* | Tropical | 6.7 | 3.9 | Longdon lab | Unknown (received 2020) |
| *Drosophila melanogaster (S)* | *melanogaster* | Temperate | 2.2 | 0.1 | Sørensen lab | Odder, Denmark (2020) |
| *Drosophila suzukii (S)* | *melanogaster* | Temperate | 2.6 | -2.8 | Colinet lab | Thorigne Fouillard, France (2016) |
| *Drosophila bunnanda (S)* | *melanogaster* | Tropical | 6.7 | 5.1 | Hoffmann lab | Australia (2008) |
| *Drosophila serrata (S)* | *melanogaster* | Tropical | 4.2 | 0.0 | David lab | Unknown (received 2014) |
| *Drosophila azteca (S)* | *obscura* | Arid | 3.6 | -7.1 | NDSSC | Andreas Canyon, California, USA |
| *Drosophila persimilis (S)* | *obscura* | Temperate | -0.7 | -6.9 | NDSSC | Victoria, British Columbia, Canada |
| *Drosophila obscura (S)* | *obscura* | Boreal | -0.6 | -8.6 | Lankinen lab | Kilpisjärvi, Finland (2020) |
| *Drosophila willistoni (S)* | *willistoni* | Tropical | 5.9 | 4.4 | Kyoto Stock Center | Guadeluope Island, France (2003) |

\*Data obtained from Kellermann*, et al.* (43).
\*\*Data obtained from Moos*, et al.* (31) except *D. willistoni* which was determined here.

Across species, we did find a significant phylogenetic relationship of [K^+^]_crit_ (Pagel’s λ = 0.960, P = 0.003) such that species within a clade are more similar to each other than those of other clades (Blomberg’s K = 0.825, P = 0.001) (**Fig. 3A**). For example, *Zaprionus* species tend to have a slightly lower [K^+^]_crit_ while species from the *Sophophora* subgenus trend towards slightly higher values. Even so, despite being tested across temperatures ranging from -11.2 to 2.5°C, [K^+^]_crit_ only varied from 35.2 to 41.9 mM across the 25 species. We also examined if the regulated baseline hemolymph [K^+^] varied across species, and despite some variance (range = 14.3 to 19.0 mM) we found no phylogenetic signal (Pagel’s λ = 0.776, *P* = 0.520) and all species had similar values overall (Blomberg’s K = 0.569, *P* = 0.052). Despite these findings, *Drosophila* species generally differ in their baseline hemolymph [K^+^] (32) and we therefore we reanalyzed data regarding the hyperkalemic response when accounting for variation in baseline [K^+^] by expressing [K^+^]_crit_ as a proportional increase above baseline [K^+^] – the relative [K^+^]_crit_ (r[K^+^]_crit_). Using this approach, we found r[K^+^]_crit_’s phylogenetic signal to disappear (Pagel’s λ = 0.171, *P* = 0.457; Blomberg’s K = 0.385, *P* = 0.396; data not depicted). Accordingly, cold stress mortality is associated with an evolutionary conserved hyperkalemic response pattern represented by a factorial increase in hemolymph [K^+^] above control (mean ± s.d. = 144 ± 15%, range = 110 to 179%), rather than coinciding with an absolute fixed [K^+^]_crit_ value. When we analyze across habitat ecotypes (**Table 1**) in phylogenetically corrected variance analyses we find no difference in either control hemolymph [K^+^] (F_3,21_ = 2.4, *P* = 0.124) or [K^+^]_crit_ (F_3,21_ = 1.2, *P* = 0.374), and by extension the r[K^+^]_crit_ is also similar across habitat ecotypes (F_3,21_ = 1.3, *P* = 0.367; not depicted) (**Fig. 3B**). To investigate this further we tested for correlations between [K^+^]_crit_ and r[K^+^]_crit_ and the species-specific biogeography or cold tolerance using phylogenetically corrected linear models but found no support for associations with either latitude (*P* = 0.843 [*t*_23_ = 0.2] and 0.480 [*t*_23_ = 0.7]), annual mean temperature (*P* = 0.844 [*t*_23_ = -0.2] and 0.338 [*t*_23_ = -1.0]), lowest temperature of the environment (*P* = 0.710 [*t*_23_ = -0.4] and 0.285 [*t*_23_ = -1.1]), 24 h LT_50_ (*P* = 0.311 [*t*_23_ = -1.0] and 0.293 [*t*_23_ = -1.1]), or CT_min_ (*P* = 0.723 [*t*_23_ = -0.4] and 0.285 [*t*_23_ = -1.1]) (see **Fig. S3**). Despite substantial interspecific variation in cold tolerance, with tolerant species surviving temperatures more than 10°C lower than sensitive species, the physiological response to equivalent cold stress was strikingly similar across species. Indeed, this conserved cold stress syndrome suggests that mortality arises from a common physiological endpoint, indicating a convergent mechanism of cold injury across the *Drosophila* genus.

To compare our findings from the *Drosophila* genus with observations from other insects orders we surveyed data from the literature for studies examining the relation between Lt_50_ and hemolymph [K^+^]. Across the few studies we found, the methods to measure both [K^+^] and injury endpoints differ, and insect species also vary substantially in their baseline (control) hemolymph [K^+^]. We therefore quantified and compared the factorial increase in hemolymph [K^+^] represented by [K^+^]_crit_ (r[K^+^]_crit_) for each species (see *Materials and Methods*) (**Fig. 3C**). In this comparative analysis we used data from five major insect orders split across holo- and hemimetabolous insects wherein the increase in hemolymph [K^+^] above baseline at Lt_50_ ranged from 111 to 194% with a mean value of 156 ± 30% (s.d.) (**Fig. 3C**, red). These values did not differ from those obtained for the 25 *Drosophila* species within this study (blue in **Fig. 3C**; Δ = 12%, CI = -65 to 46%, Monte Carlo *P* = 0.653). Although the number of species included in this broader comparison is limited by the available data, it supports that the tolerable cold stress limit is determined by the conditions leading to the same relative increase in hemolymph [K^+^] is evolutionarily conserved across a broad range of chill susceptible species representing the insect phylogeny.

### Adaptive and plastic variation in cold stress resistance

Having established a uniform physiological stress tolerance limit among insect species and acclimation groups (i.e. a uniform r[K^+^]_crit_ response) (**Fig. 2,3**), we next investigated if intra- and interspecific differences in *Drosophila* cold tolerance were associated with variation in the thermal limit that defines the onset of this cold stress syndrome. Specifically, we compared three species and five acclimation groups of the same species to record the critical homeostatic threshold temperature (T_c_) to examine to what degree these treatment groups could resist the development of the cold stress syndrome. For these experiments we exposed the flies to different constant low temperatures for 24 hours to find the warmest temperature where hemolymph [K^+^] was significantly elevated above baseline and the lowest temperature where it did increase above control. Through breakpoint analysis we then quantified T_c_, defined here as the threshold temperature that separates homeostasis (maintained baseline [K^+^]) and homeostatic deficit (hyperkalemia). We first performed this experiment in the three species tested for [K^+^]_crit_ initially and found T_c_ to range from -2.3 ± 0.7°C (*D. montana*) to 4.6 ± 0.5°C (*D. melanogaster*) and 7.7 ± 0.4°C (*D. sulfurigaster*) (**Fig. 4A**). These different levels of cold resistance match the known order of chill sensitivity, but despite clear species differences the correlation to species-specific environmental minimum temperatures did not reach statistical significance using data from only three species (*P* = 0.160, **Fig. 4A insert**). Next, we quantified T_c_ in *D. melanogaster* developmentally acclimated to the same range of temperatures as above (12 to 27°C). Here too T_c_ varied according to known degrees of cold sensitivity, ranging from 2.2 ± 0.5°C in 12°C acclimated *D. melanogaster* to 7.7 ± 0.4°C in those acclimated to 27°C and with intermediate values for acclimation groups in between (**Fig. 4B**), resulting in a strong correlation between acclimation temperature and T_c_ (*P* = 0.003, **Fig. 4B insert**). Together, these data strongly suggest that variation in *Drosophila* cold tolerance is driven by adaptive and plastic differences in the critical temperature for homeostatic regulation (T_c_). Thus, increased cold tolerance is associated with a lowered T_c_ and greater capacity to resist cold-induced disruption of ion homeostasis, a principle that may extend across chill-susceptible insects generally. Accordingly, once the species/acclimation group is exposed to temperatures below T_c_, the inability to maintain ion balance will gradually lead to the same cold stress syndrome as outlined above.

**Figure 4.**
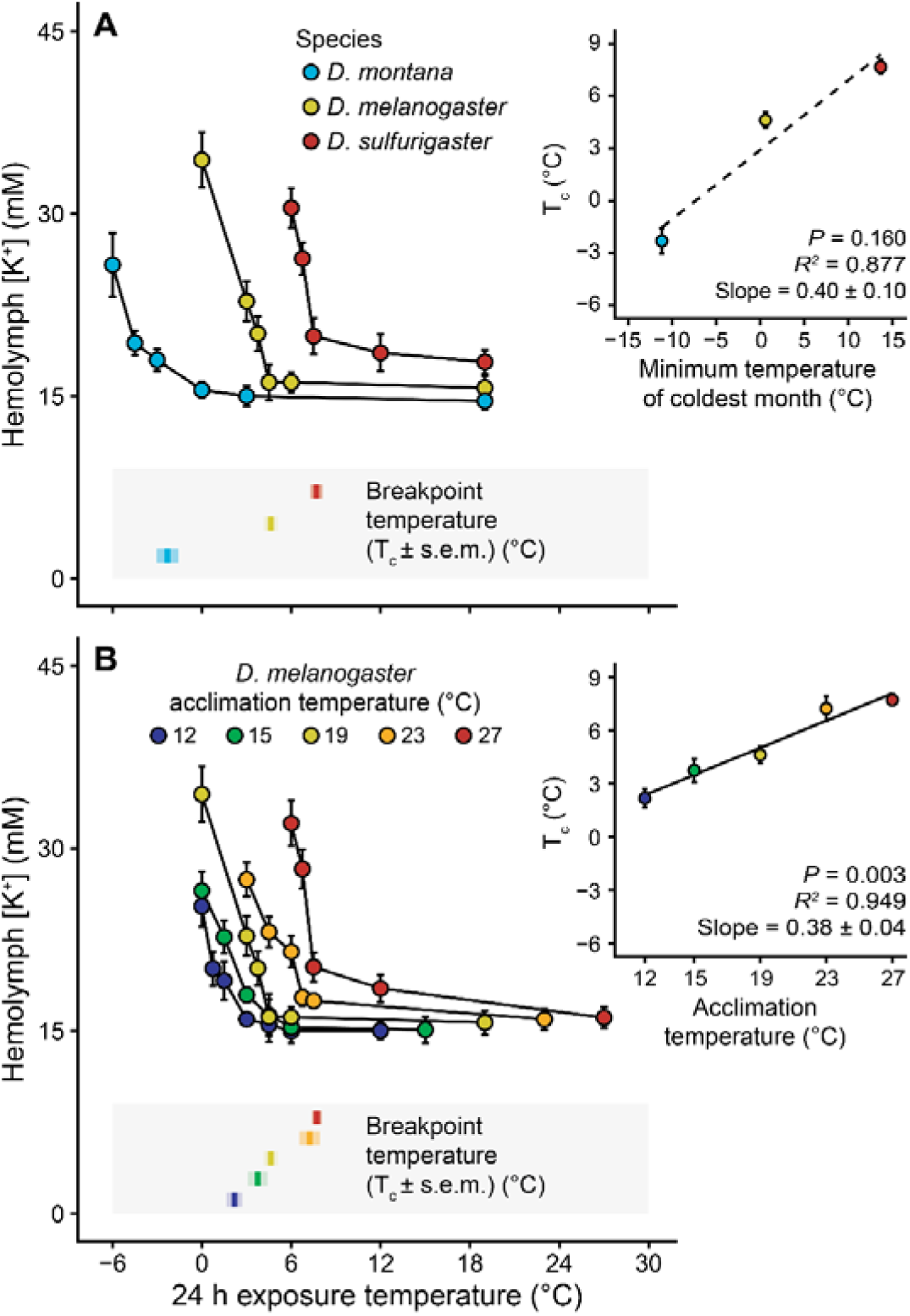
– **Variation in *Drosophila* cold sensitivity is mediated by changes in thermal resistance as shown by the systematic variation in homeostatic threshold temperature (T_c_).** (**A**) Hemolymph [K^+^] in *D. montana* (blue), D*. melanogaster* (yellow), and *D. sulfurigaster* (red) under rearing conditions (19°C) and after 24 h exposures to various low temperatures. Species-specific homeostatic threshold temperatures (T_c_) based on breakpoint analyses are indicated by color-coded means ± s.e.m in the shaded box. Insert illustrates the correlation between minimum temperature of the coldest month in the species-specific habitats (31) and the homeostatic threshold temperature (T_c_) . (**B**) Acclimation groups of *D. melanogaster* developed at five different temperatures (indicated by color). For each group, hemolymph [K^+^] was measured from flies sampled at the acclimation temperatures and in flies exposed to 24 h at various cold treatments. Acclimation group-specific T_c_ ± s.e.m. are indicated in the shaded box. Insert shows the correlation between acclimation temperature and the homeostatic threshold temperature (T_c_). Error bars not visible are obscured by the symbols. Raw data and breakpoint analyses can be found in **Fig. S4**.

## Discussion

In the present study we provide strong evidence that insect chill tolerance, the most prevalent cold tolerance strategy across insects, is determined by the species’ ability to avoid an archetypical cold stress syndrome. This syndrome develops when temperature falls below a critical threshold (T_c_), causing loss of ionoregulatory capacity resulting in a progressive rise in hemolymph [K^+^] towards a new injurious disequilibrium. Across a wide range of cold stress intensities, developmental acclimation treatments, and phylogenetically distinct *Drosophila* species, mortality consistently occurred after reaching the same degree of hyperkalemia (i.e. r[K^+^]_crit_ at ∼150% above baseline). Accordingly, across species, acclimation groups, and different cold stress intensities (Lt_50_ ranging from 1 to 25 h), insects differ primarily in the rate at which the syndrome develops and in the temperature at which it occurs, rather than in the physiological endpoint *per se*. Together, these findings indicate that cold tolerance is governed by variation in resistance to displace the critical temperature defining the onset of ionoregulatory collapse (T_c_), whereas the degree of homeostatic disruption associated with mortality (r[K^+^]_crit_) represents a strongly conserved physiological limit across insects.

### Different stress intensity, same tolerance limit

The survival duration under low temperature stress in insects, including *Drosophila*, depend directly on the stress intensity which is determined by the exposure temperature (33, 34). Accordingly, studies show that time to 50% mortality (Lt_50_) decreases predictably and exponentially with lowering of the exposure temperature used to invoke cold stress (33–35). In the present study we confirm the acceleration of cold mortality with lowering of temperature in three different species (*D. montana*, *D. melanogaster* and *D. sulfurigaster*) characterized by very different basal cold biology (**Fig. 2**). The pattern of accelerated mortality (lowered Lt_50_) with lowering of temperature found in all species, is not caused by different forms of cold stress occurring at mild vs. intense cold stress, respectively (33). Rather, we find that cold mortality is explained by the acceleration of the physiological failure cascade such that mortality is directly associated with a particular level of ionoregulatory collapse. For all three species and for all stress intensities, mortality coincides with the same critical degree of hyperkalemia ([K^+^]_crit_) (**Fig. 2A,B**). Further, when cold stress is applied in split-dose experiments combining two stress intensities for *D. melanogaster* (-3 and -7°C, respectively), we again find that progression towards the same [K^+^]_crit_ explains onset of mortality (**Fig. 2C**). These observations are consistent with existing models of insect chill injury where cold-induced hyperkalemia arises from the combination of compromised epithelial barrier integrity and reduced iono- and osmoregulatory capacity at low temperature (21). At the cellular level, hyperkalemia and low temperature synergistically induce a critical depolarization that triggers Ca^2+^ influx and intracellular Ca^2+^ overload, activating cell death pathways (22, 23), of which necrosis appears to be the dominant mechanism (24, 36). As such, hemolymph hyperkalemia remains a uniquely strong predictor of injury in cold-stressed chill sensitive insects (24).

### Increased hemolymph [K^+^] is an evolutionary conserved syndrome of insect chill tolerance

To explore putative phylogenetic differences in the critical limit of ionoregulatory failure we expanded our analysis to 25 *Drosophila* species and reconstructed their evolutionary trajectories before extending the comparison to non-*Drosophila* insects from the literature (**Fig. 3**). Although species differed slightly in their absolute [K^+^]_crit_, this variation clustered around the inferred ancestral value of ∼ 39 mM. After accounting for species-specific differences in baseline hemolymph [K^+^] (32) (r[K^+^]_crit_), a remarkably conserved tolerance limit emerged, with injury occurring at ∼ 150 % increase in hemolymph [K^+^] above baseline. This pattern persisted regardless of native habitat type (**Fig. 3B**) or latitudinal distribution (**Fig. S3**), suggesting that drosophilid flies share a common r[K^+^]_crit_ limit during cold stress. Strikingly, the pattern extends across the insect phylogeny, spanning both hemi- and holometabolous insects (**Fig. 3C**). Collectively, these findings indicate that chill susceptible insects are constrained in their cold tolerance by their ability to avoid a conserved r[K^+^]_crit_ limit that is relative to their baseline hemolymph [K^+^]. This is the first quantitative characterization of such a physiological limit and reveals an unexpected degree of conservation across insect evolution, providing insight into a previously untested hypothesis (9). The observation that the injurious limit is defined by a proportional increase in hemolymph [K^+^] (r[K^+^]_crit_) rather than an absolute change in concentration ([K^+^]_crit_) is consistent with injury occurring through membrane depolarization (23, 37), as a similar fold-increase in extracellular [K^+^] causes a similar reduction in the electrochemical gradient that sets the membrane potential (38, 39). Accordingly, this conserved limit (r[K^+^]_crit_) likely reflects a physiological threshold at which the membrane depolarization becomes sufficient to trigger activation of voltage-gated Ca^2+^ channels in insects (40, 41) causing an injurious Ca^2+^ influx that triggers cellular damage. Indeed, we here estimate the degree of depolarization caused by the combined effect of low temperature and a ∼ 150% increase in hemolymph [K^+^] to be ∼ 33 mV at Lt_50_ (see Supplementary material for calculation). This value aligns remarkably well with the degree of depolarization, ∼ 38 mV, found necessary to cause 50% cell mortality in isolated muscle preparations from chill susceptible locusts (23, 42), which have a slightly higher [K^+^]_crit_ than *Drosophila* (**Fig. 3C**).

### Inter- and intraspecific differences in Drosophila cold resistance

The conservation of a physiological limit imposed by [K^+^]_crit_ implies that adaptive and plastic variation in cold tolerance must arise from mechanisms that alter thermal resistance towards homeostatic perturbation rather than the tolerance limit ([K^+^]_crit_) itself. To investigate this, we quantified the critical homeostatic threshold temperature (T_c_) at which ion balance is disrupted by exposing flies to different low temperature treatments. Here we found inter and intraspecific differences in T_c_ that closely match established patterns of cold sensitivity among species and acclimation groups (**Fig. 4**). Specifically, species differences in T_c_ (quantified following 24 h exposures), span ∼ 10°C from *D. montana* to *D. sulfurigaster*, which aligns with their difference in 24 h LT_50_ found in our cold stress intensity experiment (∼ 11°C) and is also well aligned with species differences in CT_min_ (∼ 11°C) for these species (31, 33, 43) and in differences in literature estimates of 24 h LT_50_ (∼ 16°C). Additionally, T_c_ consistently lies ∼ 4-6°C above the 24 h LT_50_, indicating that homeostatic capacity becomes limiting well before temperatures causing mortality in this time frame. Accordingly, we find a strong association between adaptive differences in T_c_ and the habitat temperature of the species native environment (∼ 0.4°C difference in T_c_ per °C decrease in minimum environmental temperature, **Fig. 4A insert**). Although the correlation from our data using only three species does not reach statistical significance, it is entirely consistent with previous reports of adaptive changes in *Drosophila* cold tolerance limits (4, 33, 43–46). Similarly, our reported changes in T_c_ following development acclimation in *D. melanogaster* match known effects on *Drosophila* cold tolerance (∼0.4°C change per °C change in acclimation temperature, **Fig. 4B insert**) (11, 12, 46). The present study does not directly measure the mechanistic basis for the observed differences in T_c_ among or within the examined species, but both adaptive and plastic effects on *Drosophila* cold sensitivity have repeatedly been associated with enhanced ionoregulatory capacities at low temperatures that could offset the development of hemolymph hyperkalemia. For example, cold tolerant *Drosophila* are known to have superior low temperature performance of their Malpighian tubules, preserving hemolymph K^+^ removal and Na^+^ retention (47–49). Similarly, cold adaptation and cold acclimation in *Drosophila* support maintenance of hemolymph volume at low temperature (47, 50, 51), in part through increased Na^+^ reabsorption, and K^+^ excretion at the hindgut (48, 49, 52–54). These adaptations are likely facilitated by enhanced Na^+^/K^+^ ATPase (55–58) and V-type H^+^ ATPase (49, 53, 59) activity at low temperature in cold adapted/acclimated species. Finally, cold tolerant insects are typically characterized by tighter epithelial barrier function which will act to limit ion and water leak across the gut epithelium (48, 60, 61). At the cellular level maintenance of balanced ion-motive transport is likely associated with adaptive differences in phospholipid composition to optimize membrane fluidity (13, 62), continued neuroendocrine stimulation of ion transport (53), improved mitochondrial energy supply in cold adapted species (63, 64) all of which is associated with large-scale metabolomic, transcriptomic, and proteomic remodeling following cold acclimation (31, 65–67).

So far, our discussion has only concerned chill-susceptible insects as there is very limited data considering the role of ion balance for freeze-avoiding and freeze-tolerant species. Nevertheless, the limited data available suggest that these cold-tolerance strategies may share a similar [K^+^] tolerance limit. Specifically, freeze-tolerant moth larvae exposed to severe sub-zero cold exhibits mortality following a ∼ 160% increase in hemolymph [K^+^] (68), whereas freeze-tolerant fly larvae and freeze-avoidant beetle larvae are capable of stabilizing hemolymph [K^+^] below the injurious level (∼ 100% increase) and survive severe cold stress for 16 h and a week, respectively, without injury (69, 70). These observations raise the possibility that adaptation to progressively colder environments initially acts to improve resistance to ionoregulatory disruption and lower T_c._ In some cases, and often after winter acclimatization, this ability has enabled the species to tolerate such low temperature that they eventually meet a different physiological challenge – the ability to avoid or tolerate ice formation (6, 71). Further research is, however, required to determine to what extend chill tolerance, freeze avoidance, and freeze tolerance represent successive evolutionary extensions of shared homeostatic mechanisms.

## Conclusion

Collectively, our findings identify a conserved physiological limit to insect chill tolerance and provide a basis for understanding how adaptation occurs in light of this constraint. Mortality invariably occurs upon reaching a conserved threshold (r[K^+^]_crit_) representing a critical loss of homeostatic regulation. In this respect the considerable inter and intra-specific variation in cold tolerance can be explained by adaptive and plastic variation in in the threshold temperature at which ion homeostasis begins to fail (T_c_). These findings indicate that adaptation to cold proceeds primarily through mechanisms that offset the onset of physiological stress rather than through changes in the physiological limit itself. Consequently, insect physiological responses at the cellular, organ, and organismal levels should be understood through their contribution to maintenance of ion balance within homeostatic limits during low temperature exposure.

## Materials and Methods

### Animal husbandry

All *Drosophila* species populations (see **Table 1** for species information) were maintained under common garden conditions ensuring continuous development and reproduction of all species: Constant 19°C with a 22:2 light:dark cycle in 250 mL plastic bottles with ∼ 40 mL of artificial diet (grams per liter of water: 60 yeast, 40 sucrose, 30 oatmeal, 16 agar, and 12 mL nipagin and 1.2 mL acetic acid). All species tested in this study and maintained under these conditions, except *Drosophila willistoni*, had their cold tolerance tested recently (31) (temperature leading to 50% mortality in 24 h; 24 h LT_50_), and for *D. willistoni* we replicated the procedure to determine its LT_50_ (**Table 1**). Parental flies were transferred to new bottles weekly to control offspring density, and experimental flies were collected from their offspring over several days to ensure sufficient quantities and control for age. On the day of collection, flies were sorted by sex under brief CO_2_ anesthesia (< 5 min) and left for a minimum of two days before being used in experiments (72, 73).

For acclimation experiments parental *D. melanogaster* were left at 19°C to oviposit in bottles for 24 h after which the food (containing eggs) was split across new bottles and placed at acclimation temperatures of 12, 15, 19, 23, and 27°C, respectively. The flies emerging from these conditions were aged and sorted as described above, except flies acclimated to 12°C were left to recover from CO_2_ anesthesia for a minimum of four days. Because some species and cold-acclimated *D. melanogaster* (12 and 15°C) are slow to develop and because the cohorts only eclose gradually this procedure resulted in a wider age span of experimental flies, particularly from the cold acclimated flies (across experiments; approximate mean age ± s.d. = 10 ± 3 days, range = 4 to 25 days).

### Experimental overview and design

We conducted four overarching sets of experiments to investigate the relationship between ionoregulatory collapse and organismal failure in cold-stressed *Drosophila*. Across the first three experiments, flies were exposed to specific temperatures with some flies allocated to monitor survival and others were sampled to measure hemolymph [K□]. Experimental temperatures for these experiments were selected based on previously published cold tolerance data (31, 33) and pilot experiments to ensure appropriate stress intensities. The fourth experiment was designed to identify the lower critical temperature (T_c_) below which ion homeostasis collapses. For this experiment we only measure hemolymph [K^+^] sampled after a 24 h exposure time to different temperatures.

#### - Experiment 1 – Cold stress intensity

The effect of cold stress intensity was tested within three species (the chill tolerance boreal *D. montana*, the cosmopolitan temperate *D. melanogaster*, and the tropical chill sensitive *D. sulfurigaster*) where each species was exposed to three temperatures representing severe, moderate, and mild cold stress resulting in Lt_50_’s of approximately 1, 5, and 25 h. Among species and stress intensities this resulted in experimental temperatures ranging from -13.5 to 2.5°C. In addition, we tested how exposure to variable cold stress intensity (split-dose) affected ion balance. Here *D. melanogaster* were exposed sequentially to moderate (-3 °C) and severe (-7 °C) cold stress, switching between temperatures after half of the Lt_50_ duration was reached at each temperature (∼ 2.5 h at -3 °C and ∼ 27 min at -7 °C; based on the prior experiment).

#### - Experiment 2 – Plastic variation in Drosophila cold tolerance

*D. melanogaster* were reared at five developmental temperatures (see above) and subsequently exposed to acclimation-specific temperatures designed to produce a comparable moderate cold stress intensity (Lt_50_ ∼ 3-5 h). Experimental temperatures were selected based on the established relationship between developmental temperature and cold tolerance (34) and modified according to pilot experiments, resulting in exposure temperatures ranging from -5 to 2°C.

#### - Experiment 3 – Adaptive variation across Drosophila species

25 *Drosophila* species selected to represent the *Drosophila* phylogeny broadly (see **Table 1**) were used to investigate potential phylogenetic differences in the hyperkalemic stress response. All species were exposed to cold stress at temperatures designed to result in similar Lt_50_ values. The exact experimental temperatures used for each species were estimated based on data from two recent studies. In one study, the lethal temperature (LT_50_) to cause mortality after a 24 hour exposure was reported for all species used (except *D. willistoni*) (31), and in another study, the exponential relationship between tolerance duration and temperature was characterized for six species of *Drosophila* (33). These two datasets were used to approximate the temperature that should result in Lt_50_ after ∼ 3 hours, which here resulted in experimental temperatures ranging from -11.2 to 2.5°C across species.

#### - Experiment 4 – Cold stress resistance

To investigate the homeostatic threshold temperature (≈ T_c_ in **Fig. 1**) defining the lowest temperature at which a species can maintain hemolymph [K^+^] at regulated control value we prepared *Drosophila* species and *D. melanogaster* acclimation groups as in *Experiment 1* and *Experiment 2*. These flies were exposed to different low temperatures for 24 h after which hemolymph [K^+^] was measured (survival was not estimated in this experiment). Exposure temperatures for each species and acclimation group were initially chosen based on a qualified guess of the suspected threshold temperature informed by *Experiment 1* and *Experiment 2*. If [K^+^] was not elevated, we lowered temperature for the next experiment and *vice versa* until we covered at least two benign (no change) and two stressful (significant change) temperatures allowing for a high-resolution estimate of the threshold temperature.

### Lt_50_ estimates

To measure Lt_50_ in different species and under different cold exposures we exposed, for each experiment, multiple groups of flies to the assigned stressful temperature and then subsequently sampled these groups at different timepoints to measure survival allowing us to model Lt_50_. Specifically, groups of 3-7 flies were put into 200 μL pipette tips where the large opening was sealed with parafilm (a similar treatment to the flies used for measurement of hemolymph [K^+^] -see below). These flies were then put into 50 mL falcon tubes suspended in an ethylene-glycol-and-water mixture pre-cooled to the experimental temperature using a temperature-controlled cooling unit (Lauda-Brinkmann, NJ, USA). Experimental temperatures were monitored with type-K thermocouples placed in pipette tips under identical conditions. Groups of flies were subsequently sampled from the cold treatment at no less than five time points (median = 6, range = 5-12) with a median of 13 flies for each time point (range = 3-29) after which they were returned to their maintenance temperature in vials with food to allow for survival to be determined after 24 h of recovery. Flies were deemed to be alive if they were able to return to a standing position after being gently knocked over. Lt_50_ was calculated by fitting a logistic function to the survival data as follows:

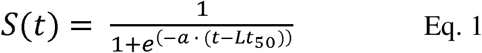

where *S* is survival expressed as the proportion of live flies, *t* is the time in hours, *e* the exponential function, and *a* and *Lt_50_* fitted variables, where *a* is the slope in the middle of the descending part of the sigmoidal curve, and *Lt_50_* its midpoint and the time point where survival reaches 50%.

### Hemolymph extraction and [K^+^] measurements

To measure hemolymph [K^+^] individual flies were loaded into 200 μL pipette tips as described above to facilitate hemolymph sampling by antennal ablation (74). To extract hemolymph pipette tips were attached to a tubing system which applied gentle air pressure from a compressor. Once the flies were pushed (head-first) into the front of the pipette, the tip was cut off, and an antenna was picked off with a pair of micro-forceps to elicit hemolymph protrusion. From here, the hemolymph droplet was placed under hydrated paraffin oil for subsequent determination of its [K^+^].

Hemolymph [K^+^] was measured with ion-selective microelectrodes (47, 75). These were fashioned from borosilicate glass capillaries (TW150-4, World Precision Instruments (WPI), Sarasota, FL, USA), pulled to a tip diameter of ∼ 1-3 μm on a Flaming-Brown micropipette puller (model P-1000, Sutter Instrument Co., Novato, CA, USA) and silanized by baking in an atmosphere of N,N-dimethylstrimethylsilylamine at 300°C for 1 h. To prepare ion-selective electrodes for measurements it was back-filled with 100 mM KCl and front-filled with K^+^-ionophore (K^+^ ionophore I cocktail B, Sigma Aldrich, St. Louis, MO, USA) to form a short K^+^-selective membrane. Subsequently, the tip of the electrode was dipped in a solution of polyvinyl chloride in tetrahydrofuran to prevent ionophore displacement, and the electrode connected to a high-impedance differential amplifier (FD223a, WPI) from which raw voltages were digitized with a MP100A A/D converter system and recorded using AcqKnowledge software (BioPack Systems, Goleta, Ca, USA). A reference electrode pulled from borosilicate glass (1B200F-4, WPI) to a long think shank and filled with 500 mM KCl was used to complete the circuit. Voltages recorded in hemolymph samples were converted into [K^+^] through reference to the voltages measured in standards with a 10-fold difference in concentration (i.e. 10 and 100 mM KCl, the difference made up with LiCl). From these voltages, hemolymph [K^+^] was calculated as follows:

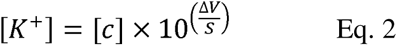

where [c] is the concentration in the lowest standard (10 mM), Δ*V* is the voltage difference between the lowest standard and the sample (in mV), and *S* is the slope of the voltage change between the two standards. The slope should follow the Nernst equation (58.2 mV) and electrodes with slopes outside of 50-62 mV were replaced. Slopes of the K^+^ selective electrodes used here were 54.6 ± 1.6 mV (mean ± s.d., based on 111 electrodes).

### Determination of hemolymph [K^+^] at Lt_50_ ([K^+^]_crit_)

Exposure to cold stress is expected to cause a progressive increase in hemolymph [K^+^] as passive ion flux exceeds the capacity for active ion transport. Because the net movement of ions is driven by the electrochemical gradient, the rate of disruption (hyperkalemia) is expected to be greatest immediately after cold exposure and to decline progressively as the gradients dissipate and the system approaches a new quasi-steady state. We therefore modelled changes in hemolymph [K^+^] using an asymptotic Michaelis-Menten-type function, because it described a rapid initial rise followed by a gradual approach to an asymptotic concentration (and not because the underlying process is assumed to follow enzyme kinetics). To estimate the degree of hemolymph hyperkalemia attained at Lt_50_ ([K^+^]_crit_) we fitted a vertically shifted Michaelis-Menten-type model to the data as follows:

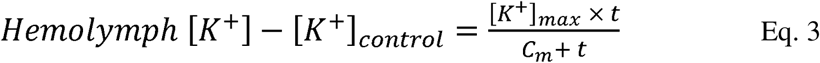

where *[K^+^]_control_* is the average control hemolymph [K^+^] for each experiment inserted as a fixed valuable to shift the model, *t* is the time in hours, and *[K^+^]_max_* and *C_m_* are fitted variables (the upper asymptote of hemolymph [K^+^] given infinite *t* and the *t* where the increase above [K^+^]_control_ is half of the *[K^+^]_max_*, respectively). From this model [K^+^]_crit_ is determined by inserting Lt_50_ in the place of *t*.

To assess the hemolymph [K^+^] attained at Lt_50_ in the split-dose additivity experiments, we used an additive model that accounted for different asymptotes and rates of increase at each temperature:

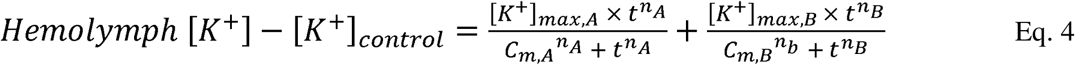

where *[K^+^]_control_* is the average control hemolymph [K^+^] inserted as a fixed value, *t* is time, *[K^+^]_max_*and *C_m_* are split into fitted variables pertaining to the first (indexed with A) or second (indexed with B) temperature, and the cooperativity exponents *n_A_* and *n_B_* for each temperature, respectively. Again [K^+^]_crit_ was determined by inserting Lt_50_ in the place of *t*. Notably, this model reduces to **Eq. 3** for single-temperature experiments as the second temperature disappears and cooperativity exponents become redundant.

Mean estimates and variation in our estimate of hemolymph [K^+^]_crit_ were obtained using a Monte Carlo resampling simulation where a random subsample of hemolymph [K^+^] data points within each dataset were procedurally removed 100 times, and the [K^+^]_crit_ subsequently estimated in the reduced datasets. Specifically, three data points were repeatedly removed from all single-temperature datasets in each simulation and the data fitted to **Eq. 3**, with five being removed from the additivity experiments and data fitted to **Eq. 4**, both representing sequential removal of ∼ 10% of data points. From these simulations we calculated mean values and standard deviations for [K^+^]_crit_ based on the 100 iterations for each dataset. Lastly, we calculated the relative degree of hyperkalemia experienced at Lt_50_ as the percentwise increase in hemolymph [K^+^] above control represented by [K^+^]_crit_, termed r[K^+^]_crit_, because *Drosophila* species can vary in their control [K^+^] (32):

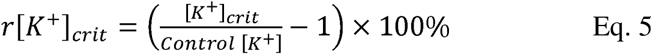

### Meta-analysis of the critical hemolymph [K^+^] in insects

Five recent studies were found to have measured hemolymph [K^+^] in relation to survival in insects in a manner that permitted comparison to the present analysis. From these five studies, raw data was extracted from four studies (22, 76–78) using WebPlotDigitizer (79) after which mortality was fitted to **Eq. 1**, and the resulting Lt_50_ used to fit hemolymph [K^+^] over time to **Eq. 3**. In the fifth study hemolymph [K^+^] was measured under control conditions and in individuals exposed to their Lt_50_ dose (80), allowing for direct calculation of the proportional increase. Notably, several studies contain data on hemolymph [K^+^] and survival in cold-stressed *Drosophila* (e.g. (49, 50, 60)), however, no study reports both Lt_50_ and temporal progression of hemolymph [K^+^] or hemolymph [K^+^] at Lt_50_. Across the five included species, control hemolymph [K^+^] varied considerably and we therefore expressed [K^+^]_crit_ as the percentwise increase above control (r[K^+^]_crit_, **Eq. 5**).

### Statistical analysis

All data analysis was performed using R software (v. 4.6.0 (81)). All analyses comparing [K^+^]_crit_ values used group-specific mean values of the Monte Carlo simulations.

#### Experiment 1

The effects of temperature and species on [K^+^]_crit_ was analyzed using a linear model with temperature as a continuous variable, species as categorical, and an interaction term. Differences in [K^+^]_crit_ between the additivity groups were assessed by calculating all possible pairwise differences between simulated [K^+^]_crit_ values from the Monte Carlo simulations (i.e. 100 x 100 = 10.000 pairs). The resulting empirical distribution of differences was used to estimate the mean difference, its 95% confidence interval, and a two-sided Monte Carlo *P*-value.

#### Experiment 2

The effect of thermal acclimation on control hemolymph [K^+^], [K^+^]_crit_, and r[K^+^]_crit_ was analyzed using linear models which included acclimation temperature as a continuous factor.

#### Experiment 3

The most recent *Drosophila* phylogeny containing all species used here (82) was used for phylogenetic analyses after being time-calibrated, which was done using the divergence time of the subgenera Drosophila and Sophophora from Suvorov et al. (83) using the ‘*ape*’ package (84). Phylogenetic signals for control [K^+^], [K^+^]_crit_, and r[K^+^]_crit_ were estimated by calculating Blomberg’s K and Pagel’s λ based on 10.000 simulations using the time-calibrated phylogeny and the phylosig() function from ‘*phytools*’ (85). Subsequently, the evolutionary trajectory of [K^+^]_crit_ was modelled by means of ancestral state reconstruction using the fastAnc() function from ‘*phytools*’. To investigate if control [K^+^] or [K^+^]_crit_ were affected by species-specific native habitat, an indicator of both cold tolerance and natural cold exposure, we categorized species based on Moos et al. (31) and tested for differences while correcting for species relatedness in the time-calibrated phylogeny with the phylANOVA() function from ‘*phytools*’ and 10.000 simulations each. Correlations between [K^+^]_crit_, r[K^+^]_crit_ and species-specific biogeography and cold tolerance were made using phylogenetically corrected linear regression with the pgls() function from ‘*caper*’ (86).

Comparison of our *Drosophila* findings to datasets available in the literature was made following the same approach as additivity groups where all empirically possible pairwise comparisons were calculated and used to estimate the mean difference, 95% confidence interval, and two-sided Monte Carlo *P*-value.

#### Experiment 4

To quantify the homeostatic threshold temperature where hemolymph [K^+^] began to deviate from control we used linear breakpoint analysis (with the segmented() function from ‘*segmented*’ (87)). Breakpoint temperatures were compared between groups using confidence intervals (calculated based on means, standard errors, and group sample sizes) and subsequently correlated to species-specific mean latitudinal distribution and acclimation temperatures, respectively, using linear regression.

Unless otherwise stated, values are presented as means ± s.e.m., except for data generated using Monte Carlo simulations (i.e. group-specific [K^+^]_crit_ values) which are presented as means ± s.d. of the simulated outputs (deviations are often obscured by points representing the mean value). Note that data for *D. melanogaster* developed at 19°C and exposed to -3°C were collected independently in *Experiments 1* and *2*. Following confirmation that the datasets were comparable, they were pooled for analyses. The level for statistical significance was 0.05 for all analyses.

## Acknowledgements

The authors would like to extend their gratitude to Ludmilla Huofková and Panalee Petcharat for their assistance during experiments.

## Author contributions

The study was conceived and designed by MKA and JO. MKA, SFJ, ESN, AS and CGB performed the experiments while MKA compiled and analyzed the data. MKA and JO wrote first draft and all authors made edits to the manuscript and approved of the final version.

## Data availability

All data collected for the present study is made available to reviewers and will be published on the *figshare* data repository upon acceptance.

